# Genotype-by-environment-by-environment interactions in the *Saccharomyces cerevisiae* transcriptomic response to alcohols and anaerobiosis

**DOI:** 10.1101/396531

**Authors:** Maria Sardi, Molly Krause, Justin Heilberger, Audrey P. Gasch

## Abstract

Next generation biofuels including longer-chain alcohols such as butanol are attractive as renewable, high-energy fuels. A barrier to microbial production of butanols is the increased toxicity compared to ethanol; however, the cellular targets and microbial defense mechanisms remain poorly understood, especially under anaerobic conditions used frequently in industry. Here we took a comparative approach to understand the response of *Saccharomyces cerevisiae* to 1-butanol, isobutanol, or ethanol, across three genetic backgrounds of varying tolerance in aerobic and anaerobic conditions. We find that strains have different growth properties and alcohol tolerances with and without oxygen availability, as well as unique and common responses to each of the three alcohols. Our results provide evidence for strain-by-alcohol-by-oxygen interactions that moderate how cells respond to alcohol stress.

**ARTICLE SUMMARY:** Efforts to promote sustainable bioenergy focus on microbial production of biofuels including butanols, which can be blended into gasoline engines and condensed for higher energy fuels. The stress of these end products can limit microbial production; yet it remains unclear how higher-energy butanols impact cell physiology compared to the well studied ethanol. This study compares the transcriptomic response to 1-butaonol, isobutanol, and ethanol in three strains of Saccharomyces cerevisiae from diverse natural populations, with and without oxygen. Results show that oxygen availability and strain background significantly impact the response to each alcohol and point to shared responses to alcohol stress.

## INTRODUCTION

The increasing interest in sustainable energy has propelled the use of genetically engineered microbes to produce fuels from non-food plant biomass. While past research has focused on ethanol production from renewable feedstock, longer-chain alcohols such as butanol and isobutanol are more attractive as next generation biofuels due to their higher energy content and compatibility with existing infrastructure for gasoline distribution. In addition, butanol isoforms are less hygroscopic and have a low freezing point, which allows for blending up to 85% with gasoline (International Energy Agency, 2011) (Durre, 2007, Fortman *et al.*, 2008). These molecules can also be modified through chemical processes to generate even more powerful sources of energy such as jet fuels (Taylor *et al.*, 2010).

Although the industrial microbe *Saccharomyces cerevisiae* does not produce significant amounts of four-chain alcohols natively, engineering *S. cerevisiae* for higher-titer butanol production is underway (Avalos *et al.*, 2013, Chen & Liao, 2016, Liu *et al.*, 2017). Production of 1-butanol in yeast is enabled by introduction of genes involved in the acetone-butanol-ethanol (ABE) clostridial fermentation, which enables the conversion of acetyl-CoA to 1-butanol (Steen *et al.*, 2008, Lian *et al.*, 2014, Generoso *et al.*, 2015, Swidah *et al.*, 2015, Schadeweg & Boles, 2016). Deletion of the alcohol dehydrogenase gene *ADH1* can also enhance native 1-butanol production (Gitter *et al.*, 2014). In contrast, production of isobutanol has been achieved by rerouting valine biosynthesis to produce isobutanol in mitochondria or the cytosol (Chen *et al.*, 2011, Brat *et al.*, 2012, Kondo *et al.*, 2012, Matsuda *et al.*, 2013, Hammer & Avalos, 2017), with highest efficiency when the whole pathway is engineered in the same cellular compartment (Avalos *et al.*, 2013, Park *et al.*, 2016). Although there have been successes in engineering *S. cerevisiae* to produce these molecules, yields are still low, making the production of next generation biofuels economically limiting at this time.

Engineering improved end-product tolerance in the host is another important consideration, since alcohol toxicity likely limits production of these fuels (Fischer *et al.*, 2008, Dunlop, 2011, Generoso *et al.*, 2015). Many studies have focused on the mechanism of inhibition caused by ethanol (Meaden *et al.*, 1999, Alexandre *et al.*, 2001, Aguilera *et al.*, 2006, Fujita *et al.*, 2006, Hu *et al.*, 2007) and identified engineering strategies that increased ethanol tolerance (Hu *et al.*, 2007, Lewis *et al.*, 2010, Swinnen *et al.*, 2012, Hubmann *et al.*, 2013, Lam *et al.*, 2014, Zyrina *et al.*, 2017). However the inhibitory mechanisms of longer-chain alcohols in *S. cerevisiae* are less well understood. Longer chain alcohols are known to cause significantly more membrane damage than ethanol (Gray & Sova, 1956, Liu & Qureshi, 2009, Huffer *et al.*, 2011), disrupting pH balance and inhibiting important membrane proteins such as ATPases and glucose transporters (Paterson *et al.*, 1972, Grisham & Barnett, 1973, Ingram, 1976, Bowles & Ellefson, 1985, Gonzalez-Ramos *et al.*, 2013). In addition, these alcohols are known to induce the accumulation of misfolded proteins (Dunlop, 2011, Ghiaci *et al.*, 2013, Gonzalez-Ramos *et al.*, 2013, Navarro-Tapia *et al.*, 2016). It is largely unclear how alcohol tolerance is affected under anaerobic conditions, which is an important consideration given that many industrial processes are performed anaerobically. Both anaerobiosis and alcohols affect membranes, but in different ways. The absence of oxygen prevents the production of sterols and unsaturated fatty acids and thus alters membrane composition and fluidity (Wilcox *et al.*, 2002, Rosenfeld & Beauvoit, 2003), while alcohols target membrane integrity directly (Ingram, 1976, Ingram, 1986, Mishra & Prasad, 1989, Alexandre *et al.*, 1994). In turn, membrane composition influences alcohol tolerance (Mannazzu *et al.*, 2008, Henderson *et al.*, 2011, Henderson *et al.*, 2013, Archana *et al.*, 2015), although the optimal membrane composition and the mechanisms of tolerance remain unclear (Huffer *et al.*, 2011, Henderson & Block, 2014). Other studies have linked respiration, protein folding, and protein degradation to aerobic butanol tolerance (Ghiaci *et al.*, 2013, Gonzalez-Ramos *et al.*, 2013, Crook *et al.*, 2016). Exploring how hypoxia modifies the inhibitory effects of alcohols has important industrial applications. But an additional important consideration is that different strains may vary in their responses and the mechanisms they use to survive anaerobic stresses.

Here, we used genetic and genomic approaches to explore the differences in alcohol response in the presence and absence of oxygen, across multiple genetic backgrounds of yeast. Our past strategy compared stress responses across natural strains with varying tolerance to identify primary targets of industrial stressors and defense mechanisms employed by tolerant strains (Lewis *et al.*, 2010, Sardi *et al.*, 2016, Sardi *et al.*, 2018). Here we compared the transcriptome responses to 1-butanol, isobutanol, and ethanol in both aerobic and anaerobic conditions, across three strain backgrounds with varying tolerances, using RNA sequencing (RNA-seq). The strains showed significant differences in their response to anaerobic growth alone and in their tolerance and response to alcohols under aerobic and anaerobic conditions. Comparing and contrasting across alcohols and aerobic conditions revealed unique responses to ethanol versus longer-chain alcohols and synergistic effects between alcohols and oxygen depletion. This study therefore revealed important genotype by environment by environment interactions that affect the stress response and alcohol tolerance under different conditions. Together, these results expand our knowledge of alcohol responses for future engineering strategies while presenting important information about how environments and genotype interact in a stress response.

## METHODS

### Strains and growth conditions

Strains used in this study and their phenotypes are listed in Table S1. Three strains were selected for more detailed analysis: West African strain NCYC3290 (WA), mosaic isolate from Illinois, USA IL01 (courtesy Justin Fay) and a mosaic strain isolated from papaya, Y7568 (PAP). The concentrations of ethanol, butanol, and isobutanol were selected as the maximal tested doses that allowed cell growth in all three strains. Unless otherwise noted, cells were grown in shaking flasks at 30°C. Rich lab medium (YPD: 1% yeast extract, 2% peptone, 2% dextrose) was used as the base medium for all growth conditions. Anaerobic growth was performed in an anaerobic chamber maintained at O_2_ < 25 PPM, using culture flasks with a stir bar to maintain cell suspension. Phenotypes for Figure 1B-C were measured using 1% 1-butanol, 1.2% isobutanol, and 5% ethanol.

**Figure 1.**
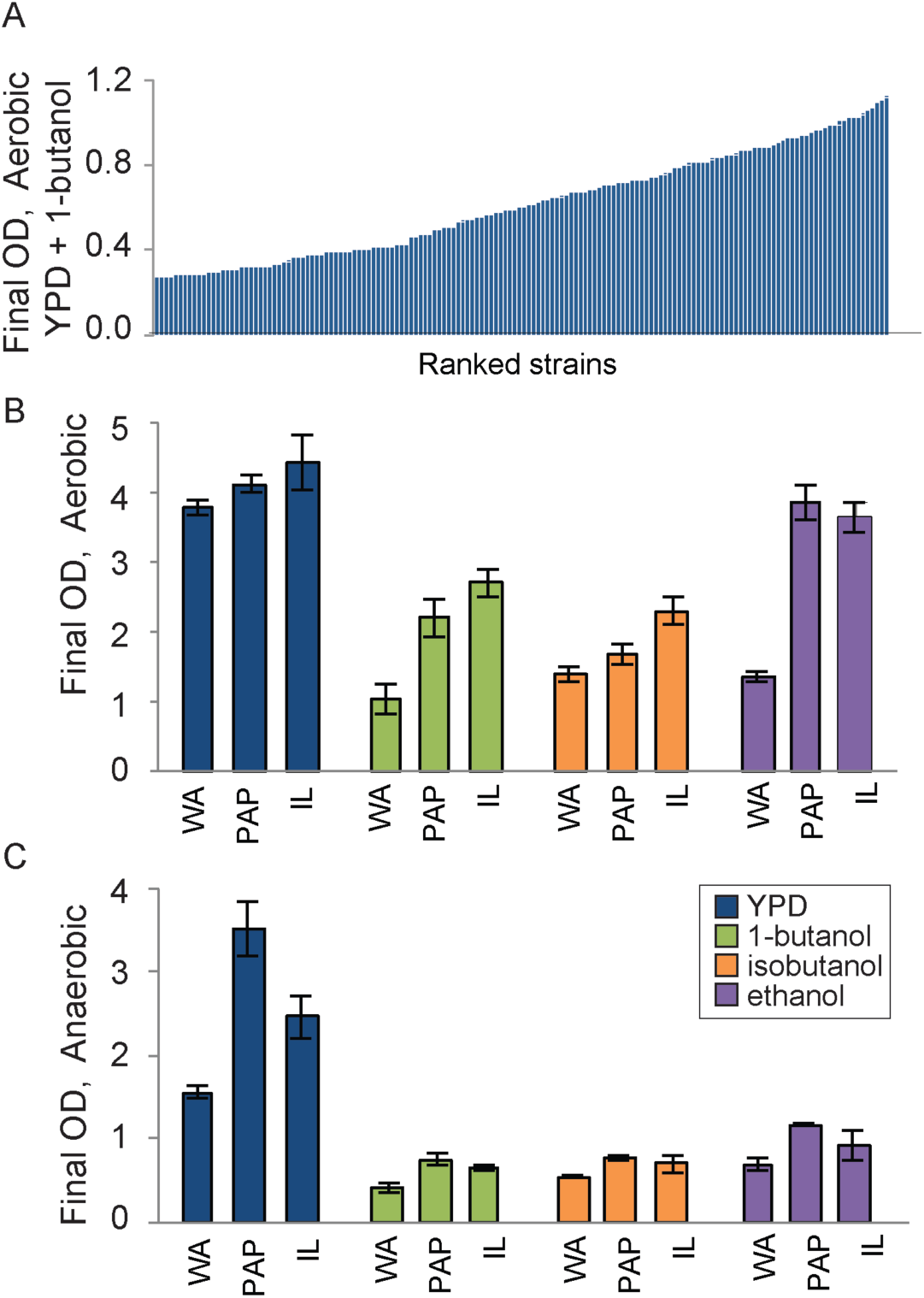
Strain-specific tolerance of alcohols varies with oxygen. (A) Representative final OD_600_ measurements of 165 strains grown in YPD with 2% butanol. Strains are sorted by final OD (Table S1). (B-C) West African strain NCY3290 (WA), mosaic strains Y7568 (PAP) and IL01 (IL) were grown in YPD with and without added alcohols. Final OD_600_ of strains grown (B) aerobically 10 h or (C) anaerobically for 11 h in YPD, YPD + 1% butanol, 1.2% isobutanol, or 5% ethanol. Data represent the average and standard deviation of 3 replicates.

### Phenotyping

High-throughput phenotyping of the 165-strain *Saccharomyces cerevisiae* collection was performed in 96-well plates (NUNC, Thermo Scientific, Rockford, IL). Briefly, 10 µl of thawed frozen stock of cells was used to inoculate a 96-well plate containing 190 µl of YPD media. Plates were sealed with breathable tape (AeraSeal, Sigma, St. Louis, MO), covered with a lid and incubated at 30°C while shaking for 24 hours, at which time a new subculture was generated by inoculating 190 µl of YPD with 10 µl of the previous culture and grown to log phase for 6 hours. 10 µl of this log-phase culture was inoculated into 190 µl of YPD + 2% butanol, plates were then sealed with an aluminum foil seal (Cryostuff, Vienna, VA, USA) to minimize butanol evaporation, grown with shaking for 24 hours, followed by measurement of the final OD_600_ using the Tecan M200 Pro microplate reader (Tecan Systems, Inc., San Jose, CA). The average of four biological replicates was calculated to represent tolerance to butanol. Single-strain phenotyping was performed in shaker flasks at 30C under defined media conditions.

### Oxygen utilization rate measurement

Strains were grown in rich lab media (YPD) for three generations and kept in log phase before oxygen measurements. A microoptode oxygen probe (Unisense, Denmark) was used to measure dissolved oxygen over time, measuring every 15 seconds for 3 minutes. Each measurement was normalized to underlying OD_600_ and oxygen consumption was taken as the slope of measured oxygen over time.

### Transcriptome profiling

Strains were grown at 30°C with shaking to mid-log phase for seven generations in YPD. The culture was then used to inoculate YPD, YPD + 0.8% 1-butanol, YPD + 1% isobutanol, YPD + 4% ethanol (or YPD + 3% ethanol for anaerobic conditions, since WA did not grow well anaerobically on 4% ethanol), grown for three generations and collected during log phase by centrifugation. Collections were performed in duplicate on different days, such that replicate pairs could be analyzed using paired statistics. Analyzing all 12 samples together with a multi-factorial linear model provided additional statistical power (see below). RNA was processed as previously described (Sardi et al. 2016). Briefly, RNA was extracted by hot phenol lysis (Gasch, 2002). Total RNA was DNase-treated at 37°C for 30 min with TURBO DNase (Life Technologies, Carlesbad, CA), followed by RNA precipitation at -20°C in 2.5M LiCl for 30 min. rRNA depletion and library generation was constructed by the University of Wisconsin-Madison Biotechnology Center, via the TrueSeq® Stranded Total RNA Sample Preparation Guide (Rev.C) using the Illumina TrueSeq® Stranded Total RNA (yeast) kit (Illumina Inc., San Diego, California, USA). Libraries were standardized to 2 µM. Cluster generation was performed using standard cluster kilts (vs) and the Illumina cluster station. Single-end 100 bps reads were generated on an Illumina HiSeq2500 sequences. All sequencing data are available in the NIH GEO database under accession number GSE118069.

Reads were processed with Trimmomatic (Bolger *et al.*, 2014) and mapped to reference genome S288c (NC_001133, version 64 (Engel *et al.*, 2014)) using bwa mem (Li, 2013) with default settings. HTseq version 5.5 (Anders *et al.*, 2015) was used to sum read counts for each gene. Differential expression analysis was performed using the program edgeR v.3.8.6. (Robinson *et al.*, 2010) using a linear model to simultaneously analyze all twelve samples across two environments, with strain background and media type as factors and replicate samples paired. Data were normalized for visualization using the reads per kilobase per million mapped reads (RPKM) method. Hierarchical clustering analysis was performed using the R package Mclust (Scrucca *et al.*, 2016) using model ‘VII’. Visualization was performed with the program Java Treeview (http://jtreeview.sourceforge.net/) (Saldanha, 2004). Where indicated, data were clustered based on the RPKM values normalized to the mean value for each transcript across all strains (‘mean centered’). Functional enrichment analysis was performed by in-house scripts for the hypergeometric test using four different datasets previously defined (Chasman *et al.*, 2014) or using FunSpec (Robinson *et al.*, 2002, Boyle *et al.*, 2004).

## RESULTS

### Alcohol tolerance varies across *Saccharomyces cerevisiae* strains and with oxygen availability

To characterize variation in alcohol tolerance, we first surveyed 165 strains of *Saccharomyces cerevisiae* collected from a variety of geographical locations and ecological niches for their ability of cells to grow in 1-butanol (Table S1, see Methods). 1-butanol tolerance is clearly a quantitative trait, and we observed a wide range of tolerances spanning four-fold variation in final cell density of the most sensitive and tolerant strains (Fig 1A). Based on these phenotypes, we chose three strains with different degrees of tolerance for more detailed investigation: a West African strain isolated from bili wine, NCYC3290 (WA), a mosaic strain isolated from a rotten papaya Y7568 (PAP) and a mosaic strain isolated from soil IL01 (IL) (Table S1).

We measured growth of our selected strains in rich laboratory medium (YPD) and in medium supplemented with 1% 1-butanol, 1.2 % isobutanol, and 5% ethanol, under aerobic (Fig 1B) and anaerobic conditions (Fig 1C). Interestingly, anaerobiosis affected strains in different ways. IL01 followed by PAP were most tolerant to butanols, whereas both IL01 and PAP showed similar aerobic tolerance to ethanol. WA was the most sensitive to all alcohols but showed particularly poor growth in ethanol (Fig 1A). However, the strains showed different phenotypes under anaerobic growth: in the absence of alcohols, PAP was the fastest growing strain and grew to nearly the same final cell density as in aerobic conditions, unlike the other strains that grew slower anaerobically. All of the strains were more sensitive to alcohols anaerobically, and the differences in tolerance to this dose of alcohol were minimal under anaerobic conditions (although WA was unable to survive higher doses of ethanol anaerobically). Thus, the results show an interesting genotype (strain) by environment (oxygen availability) by environment (alcohol exposure) interaction, such that strains show significant differences in ranked tolerance depending on the alcohol and the oxygen status.

### Strain-specific transcriptomic responses to anaerobic conditions

The condition-specific differences in strain performance provided an opportunity to investigate differences in tolerance mechanisms, with and without available oxygen. We first analyzed transcriptomes of the three strains grown in rich YPD medium with and without oxygen. There were 474 differentially expressed genes (FDR <1%) across the three strains grown aerobically (Fig 2A), and hierarchical clustering revealed groups of functionally related genes (Dataset 1). Interestingly, one cluster that was expressed higher in the fastest growing IL01 was significantly enriched for genes involved in aerobic respiration (Fig 2A), raising the possibility that IL01 relies more on respiration than the other studied strains even in the presence of glucose. To test this, we measured oxygen consumption during aerobic growth on rich medium. Indeed, IL01 consumed oxygen at a significantly faster rate compared to WA and IL01 (Fig 2B), which could contribute both to its faster growth rate and improved alcohol tolerance specifically in aerobic conditions (see Discussion).

**Figure 2.**
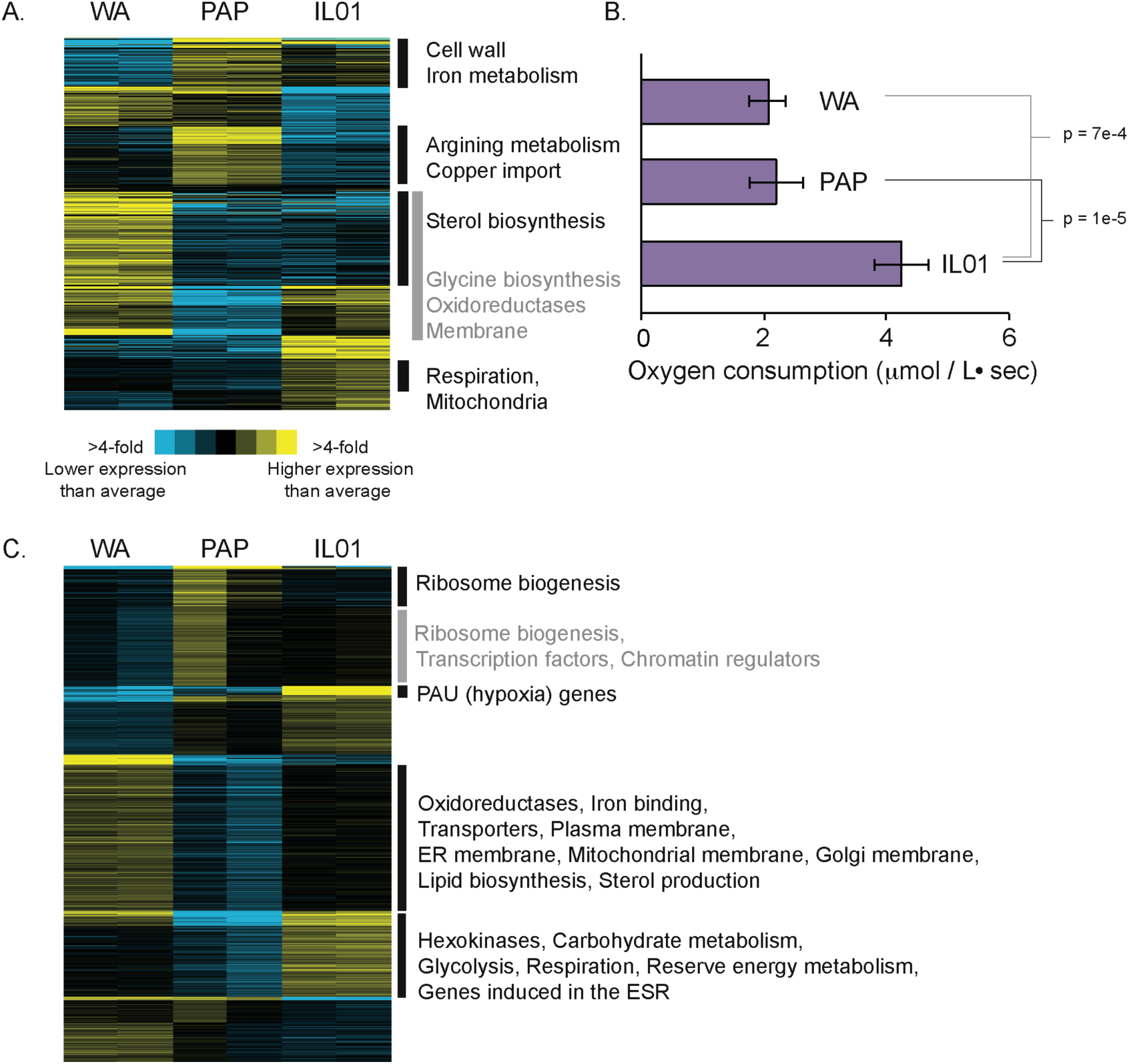
Strain-specific transcriptomic differences vary with oxygen. (A) Genes (rows) were clustered based on the mean-centered log_2_ RPKM. Shown are 474 genes whose expression was significant different (FDR < 1%) in WA, PAP, or IL01 compared to the mean expression of the three strains. (B) Rates of oxygen consumption during aerobic YPD growth. Asterisk indicates significant differences compared to the other strains. (C) Shown are 1,923 differentially expressed in each strain compared to the mean (FDR < 1%), as described in (A). Functional enrichments (p<1e-5) are listed for each group; grey boxes are used simply for demarcation.

Next, we investigated transcriptome differences under anaerobic conditions in the absence of added alcohols: the number of differentially expressed genes across strains was over three times greater than under aerobic conditions at 1,923 genes (FDR < 1%, Fig 2C, Dataset 2). Interestingly, it was the slowest growing strain WA that showed high expression of many genes, enriched for those related to membrane, ergosterol, and fatty-acid synthesis (Fig 2C). This was intriguing, because WA was the slowest growing strain under anaerobic conditions (predicted to impact membrane fluidity) and the most sensitive to membrane-targeting alcohols (see Discussion). IL01 showed significantly higher expression of genes related to energy production, mobilization of energy reserves, gene induced in the environmental stress response, and PAU genes that are induced as part of the hypoxic response (Rachidi *et al.*, 2000, Luo & van Vuuren, 2009), whereas the fastest anaerobic grower PAP displayed the lowest expression of these genes (see Discussion).

### Transcriptome responses to aerobic alcohol exposure implicate common and alcohol-specific responses

To investigate mechanisms of alcohol tolerance, we next compared the transcriptome responses to 0.8% 1-butanol, 1% isobutanol, and 4% ethanol in strains growing under aerobic conditions. We used a linear model to identify genes with whose expression was responsive to alcohol independent of strain background and genes affected by a strain-by-media (*i.e.* gene-by-environment) interaction (see Methods). We identified hundreds of genes whose expression responded to each alcohol, and many genes with strain-by-alcohol interactions (Fig 3). We hierarchically clustered the combined set of 776 genes that responded to any of the three alcohols (including those with strain-specific alcohol responses, Fig 3A, Dataset 3).

**Fig 3.**
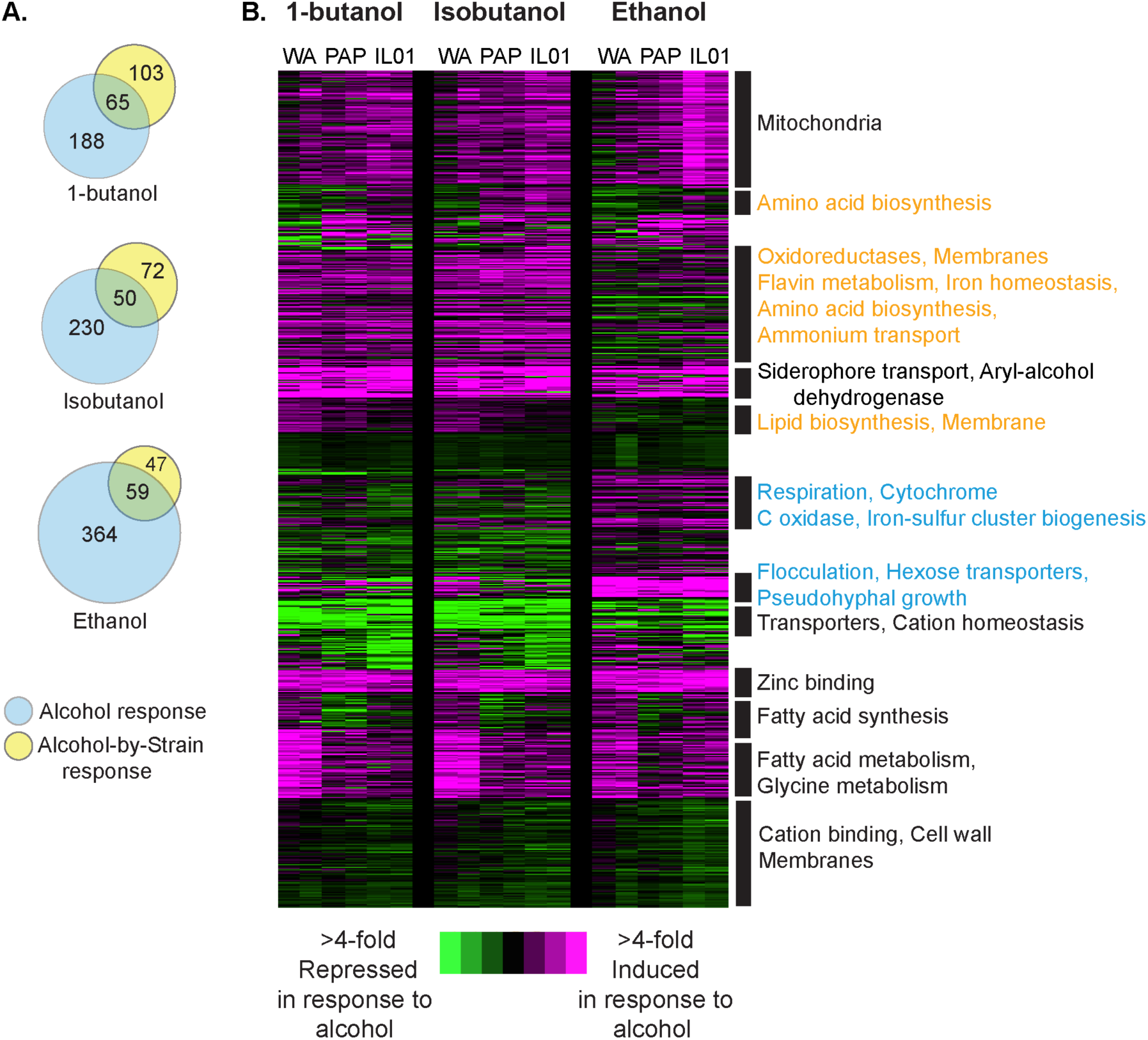
Expression responses to butanol, isobutanol, and ethanol in aerobic conditions. (A) Venn diagrams represent the number of genes whose expression responds to alcohol independent of strain and genes with a strain-specific response, for each alcohol. (B) Hierarchical clustering of 776 genes identified as differentially expressed in response to 0.8% butanol, 1% isobutanol, or 4% ethanol compared to YPD, in cells growing aerobically. Genes were clustered based on the log_2_(fold change) in expression in each strain (WA, PAP, IL) grown in the indicated alcohol versus YPD, in biological duplicate. Expression changes are colored to indicate induciton (pink) and repression (green) according to the key. Functional categories enriched in each cluster (p<1e-4, FunSpec) are shown to the right of each cluster. Responses specific to ethanol are highlighted in blue text and those specific to butanols are highlighted in orange text.

Many of the expression responses were similar across strains and regardless of alcohol identity. This included induction of genes involved in mitochondrial functions, siderophore transport, aryl-alcohol dehydrogenases, and zinc binding, and repression of genes encoding transporters and many other membrane proteins (Fig 3B). In contrast, there were also several responses unique to 1-butanol and isobutanol compared to ethanol. Butanol isoforms induced a different set of genes linked to membrane synthesis, as well as genes associated with oxidoreduction, iron homeostasis, amino-acid biosynthesis, and ammonium transport. Interestingly, ethanol uniquely triggered in all three strains the induction of genes involved in flocculation, hexose transport, pseudohyphal growth, and respiration, whereas butanol isoforms did not (Fig 3B). These responses are also seen in cells undergoing filamentous growth, and ethanol is known to trigger the response in some strains (Lorenz *et al.*, 2000). There were also several aspects of the responses that were different across strains. Most notably, WA showed much stronger induction of lipid and fatty-acid biosynthesis genes, especially in response to butanols, again suggesting that this strain may experience specific defects in membrane function (see Discussion).

### Strain-specific responses to alcohol are different anaerobically

Most industrial fermentations are anaerobic, however the majority of studies analyzing alcohol toxicity are based on aerobic conditions. We therefore analyzed the transcriptome response to each alcohol anaerobically, comparing the response to 0.8% 1-butanol, 1% isobutanol, and 3% ethanol to anaerobic YPD growth. Pooled together, the analysis identified 2,075 genes that were differentially expressed in response to alcohols in one or more strains (Dataset 4). All three alcohols were more inhibitory in anaerobic conditions (see Fig 1), and we correspondingly observed 2.6-fold more genes differentially regulated in response to alcohols under anaerobic versus aerobic conditions (Fig 4). Many of the responses were common to all three alcohols; however, we observed one gene cluster with a specific response to butanols (Fig 4B, orange text). This group of induced genes was enriched for genes related to membranes, mitochondrial function, and Gcn4 targets involved in amino acid biosynthesis. These genes were either not strongly induced or in the case of WA were strongly repressed in response to ethanol. WA had other unique responses to ethanol and likely contributed to the large number of strain-by-ethanol responses (Fig 4A). In contrast, the anaerobic response to butanols was largely similar across the three strains.

**Fig 4.**
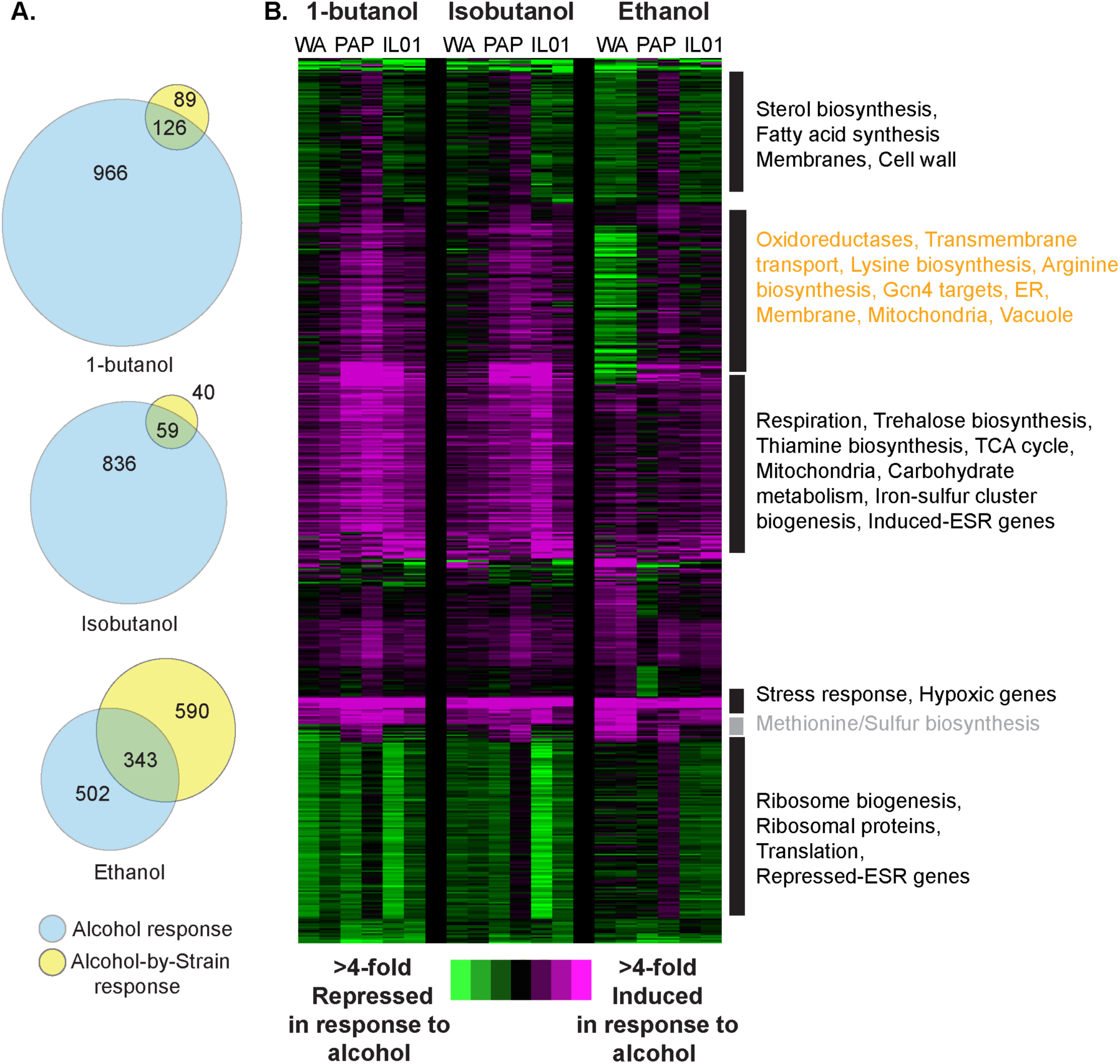
Expression responses to butanol, isobutanol, and ethanol under anaerobic conditions. (A) Venn diagrams represent the number of genes whose expression responds to alcohol independent of strain and genes with a strain-specific response, for each alcohol treatment administred anaerobicaly. (B) Hierarchical clustering of 2,075 genes differentially expressed in response to 0.8% butanol, 1% isobutanol, or 3% ethanol compared to YPD, under anaerobic conditions, as described in Figure 3.

To better understand the differences in alcohol responses, independent of strain-specific responses, we compared the alcohol-responsive genes identified by each linear model (FDR < 1%) when cells were grown with and without oxygen (Fig 5) and investigated functional enrichment among genes in each set. This provides a complementary analysis to the clustering described above. Anaerobically, all three alcohols triggered the induction of genes related to energy metabolism. These include genes involved in respiration, TCA cycle, energy reserve metabolism, and/or the stress response, and together suggests energy limitation during anaerobic alcohol defense (see Discussion). Aerobically, butanols induced genes involved in iron homeostasis, oxidoreduction, and amino-acid biosynthesis, with isobutanol producing a broader effect across Gcn4 targets as well as other genes related to lysine, arginine, and purine biosynthesis (see also Fig 4). Butanols triggered reduced expression of respiration genes during aerobic growth but induced expression anaerobically – this was in contrast to ethanol that led to the induction of respiration genes both aerobically and anaerobically (Fig 3-5). In at least two of the three strains, ethanol also triggered reduced expression of genes encoding plasma membrane and cell surface markers (Fig 5, 4B) that were not strongly triggered by butanols despite their higher toxicity. These results suggest cellular targets of the different alcohols and mechanisms cells use to respond.

**Fig 5.**
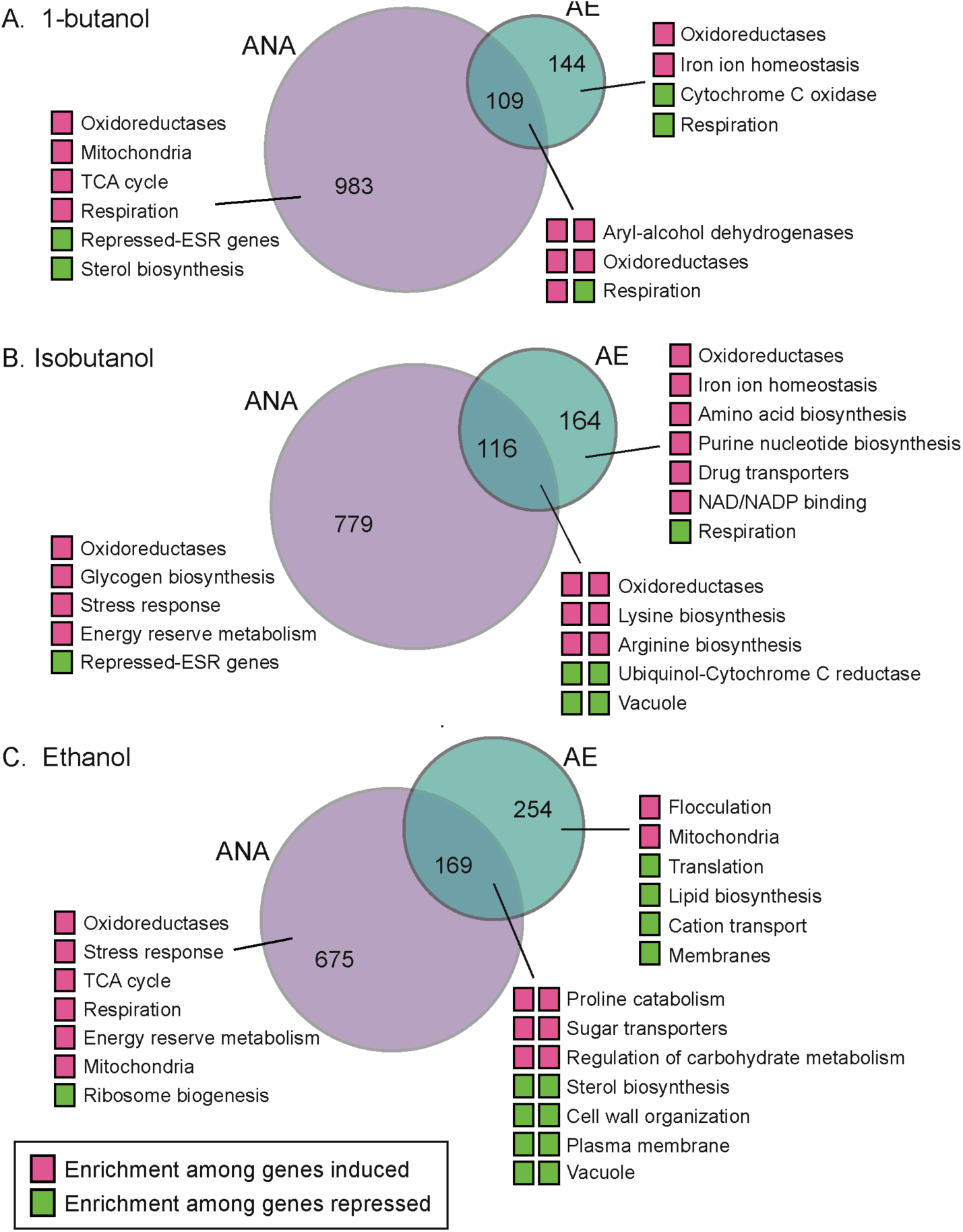
Comparison of aerobic and anaerobic alcohol responses. Each Venn diagram compares genes differentially expressed (FDR < 1%) under aerobic (AE), anaerobic (ANA), or both conditions for (A) 1-butanol, (B) isobutanol, and (C) ethanol. Colored boxes represent genes induced (magenta) or repressed (green) under those conditions; for genes in the overlap, the left box indicates anaerobic conditions and the right box represents aerobic conditions. Functional categories enriched for each gene group (p<1e-4, FunSpec) are shown.

## DISCUSSION

With the pursuit of more energy-intensive renewable fuels, recent efforts have been focused on microbial production of butanol isoforms using both synthetic and metabolic engineering strategies (Durre, 2011, Buijs *et al.*, 2013). While butanols are expected to target similar cellular processes as the better studied ethanol, the unique features of butanol stress responses are not well understood. Our comparative transcriptomic response has generated new insights into how genetics and environment influence the cellular response to these alcohols.

While we set out to study alcohol responses, an important result of our study is that different strains of *S. cerevisiae* respond differently to anaerobic growth in the absence of added stress. We were surprised to find growth properties and relative alcohol sensitivities would differ across strains depending on oxygen availability. IL01 grew on par with the PAP strain aerobically but grew significantly slower anaerobically (Fig 1); that this strain respires at a higher rate than other strains aerobically (Fig 2B) suggests that it relies more on respiration for energy generation, and hence suffers a disadvantage when respiration is blocked by anaerobiosis. In the absence of added alcohols, the WA strain showed uniquely high expression of genes enriched for membrane functions, sterol synthesis, and fatty acid production, both aerobically and anaerobically. Many of these genes were repressed in WA responding to alcohols, especially ethanol anaerobically (Fig 4). Since membrane integrity and fluidity are primary targets of alcohols, we suggest that the increased sensitivity of the WA strain may be due to an underlying difference in membrane composition or the ability to maintain it. These results highlight the utility of incorporating a comparative genomic approach to studying the response to industrial stresses that can vary significantly across strains.

Our results also point to commonalities and unique features in the responses to different alcohols across multiple strains. All strains were more sensitive to alcohols anaerobically, and consequently all strains showed more alcohol-responsive expression changes under these conditions. The common induction of genes related to energy metabolism is consistent with the idea of energy shortages during anaerobic alcohol defense. Genes involved in respiration and alternate energy mobilization are commonly induced during stress even if the absence of oxygen (Gasch *et al.*, 2000, Lahtvee *et al.*, 2016, Malina *et al.*, 2018). Furthermore, up-regulation of respiration proteins was associated with evolved 1-butanol tolerance in yeast (Ghiaci *et al.*, 2013). Yet, here we found that both 1-butanol and isobutanol treatment led to reduced aerobic expression of respiration genes. The reason for this is unclear, but could be related to other mitochondrial processes influenced by butanol response, including NADH/NAD+ rebalancing or iron-sulfur cluster generation.

Prior investigation of butanol responses in yeast and bacteria pointed to signatures of protein misfolding and oxidative stress, involving protein-folding chaperones, proteasome-dependent protein degradation, and redox rebalancing (Rutherford *et al.*, 2010, Ghiaci *et al.*, 2013, Gonzalez-Ramos *et al.*, 2013, Crook *et al.*, 2016). We observed multiple protein folding chaperones induced by butanols under anaerobic stress, but we did not see induction of proteasome genes as a group (Datasets 3, 4). Many oxidoreductases, including alcohol and aldehyde dehydrogenases (including *ADH5, ALD5, GDH1, MDH2, DLD1*) were induced uniquely by butanols, while others that are part of the yeast Environmental Stress Response (Gasch *et al.*, 2000) were induced by all alcohols. Several of these oxidoreductases may represent specific detoxification mechanisms against butanols. Finally, 1-butanol and especially isobutanol led to the induction of many genes involved in amino acid biosynthesis, including biosynthetic genes regulated by the transcription factor Gcn4. Gcn4 can be activated by amino acid starvation but also uncharged tRNAs or translation defects (Hinnebusch, 2005). It is unclear why the response is induced here, but alcohols are known to inhibit translation elongation (Ashe *et al.*, 2001, Haft *et al.*, 2014). It is also possible that the response is triggered by isobutanol itself, produced as a byproduct of valine biosynthesis; however, we saw no notable expression changes of ILV genes involved in that pathway.

Our results also raise important considerations about genotype-phenotype relationships and how they change in different environments. Both growth properties and gene expression varied dependent on strains as well as the presence of oxygen, alcohols, or both conditions, revealing important strain-by-oxygen-by-alcohol interactions. While genotype-by-environment effects are well appreciated in quantitative genetics, combinatorial effects of more complex environmental changes are generally less well studied. Our results set the stage for further investigation of genotype-by-environment-by-environment changes and how they can be leveraged for understanding and engineering industrially relevant traits.

## ACKNOWLEDGEMENTS

We thank M. Place for bioinformatic assistance. MS was supported by a This work was supported by a predoctoral fellowship from the NSF. Work was funded by the Great Lakes Bioenergy Research Center, U.S. Department of Energy, Office of Science, Office of Biological and Environmental Research under Award Numbers DE-SC0018409 and DE-FC02-07ER64494.

## SUPPLEMENTARY TABLES AND DATASETS

**Table S1:** Strains used in this study and their final OD600 in 1-butanol screen.

**Dataset 1:** Gene mean-centered log2(RPKM) expression data in strains grown aerobically on YPD medium, as shown in Figure 2A. Tab 1 contains column legend.

**Dataset 2:** Gene mean-centered log2(RPKM) expression data in strains grown anaerobically on YPD medium, as shown in Figure 2B. Tab 1 contains column legend.

**Dataset 3:** Log2(fold change) in expression comparing each strain grown in medium with the designated alcohol concentration (see Methods) versus cells grown in YPD, both grown in aerobic conditions, as shown in Figure 3.

**Dataset 4:** Log2(fold change) in expression comparing each strain grown in medium with the designated alcohol concentration (see Methods) versus cells grown in YPD, both grown anaerobically, as shown in Figure 4.

